# Enhanced protein synthesis is a defining requirement for neonatal B cell development

**DOI:** 10.1101/2022.11.11.515795

**Authors:** Hugo Åkerstrand, Elena Boldrin, Giorgia Montano, Stijn Vanhee, Karin Olsson, Niklas Krausse, Stefano Vergani, Maciej Cieśla, Cristian Bellodi, Joan Yuan

## Abstract

The LIN28B RNA binding protein exhibits a ontogenically restricted expression pattern and is a key molecular regulator of fetal and neonatal B lymphopoiesis. It enhances the positive selection of CD5+ immature B cells early in life through amplifying the CD19/PI3K/c-MYC pathway and is sufficient to reinitiate self-reactive B-1a cell output when ectopically expressed in the adult. In this study, interactome analysis in primary B cell precursors showed direct binding by LIN28B to numerous ribosomal protein transcripts, consistent with a regulatory role in cellular protein synthesis. Induction of LIN28B expression in the adult setting is sufficient to promote enhanced protein synthesis during the small Pre-B and immature B cell stages, but not during the Pro-B cell stage. This stage dependent effect was dictated by IL-7 mediated signaling, which masked the impact of LIN28B through an overpowering stimulation on the c-MYC / protein synthesis axis in Pro-B cells. Importantly, elevated protein synthesis was a distinguishing feature between neonatal and adult B cell development that was critically supported by endogenous *Lin28b* expression early in life. Finally, we used a ribosomal hypomorphic mouse model to demonstrate that subdued protein synthesis is specifically detrimental for neonatal B lymphopoiesis and the output of B-1a cells, without affecting B cell development in the adult. Taken together, we identify elevated protein synthesis as a defining requirement for early-life B cell development that critically depends on *Lin28b*. Our findings offer new mechanistic insights into the layered formation of the complex adult B cell repertoire.

## Introduction

B-cell development is centered around alternating stages of cellular proliferation and V(D)J recombination at the immunoglobulin heavy and light chain loci, with the goal of producing a diverse yet self-tolerant B cell repertoire. During the Pro-B cell stage, productive IgH rearrangement results in the expression of a pre-BCR complex and differentiation into the large Pre-B cell stage. The latter is marked by a transient burst of clonal expansion. Cell cycle exit is accompanied by the initiation of light chain recombination and entry into the small Pre-B cell stage. IL-7 signaling is an essential trophic factor during the Pro- and large Pre-B cell stages, promoting cell survival, proliferation, and differentiation, through distinct effects mediated by the Stat5 and mTORC1/c-MYC signaling axes (1–2). IL-7 signaling wanes as large Pre-B cells move away from IL-7 producing stromal cells in vivo and its attenuation is required for the developmental progression into the small Pre-B cell stage and the onset of IgK light chain recombination (3). Generation of a productive immunoglobulin heavy and light chain complex leads to membrane IgM expression and developmental progression into the immature B (Imm-B) cell stage. Here, cells carrying innocuous BCRs are allowed to mature while self-reactive specificities are subject to central tolerance mechanisms including receptor editing and apoptosis to remove potentially harmful recognition of “self’ (4). Progression through these stages is accompanied by dramatic fluctuations in bioenergetic states. Although metabolic capacity has been proposed to modulate the central tolerance checkpoint (5), this theory has not been empirically proven.

Much of what we know about B cell development comes from studies in adult mice, which differs significantly from fetal and neonatal life. For example, while B cell development in adult bone marrow strictly requires IL-7 mediated signaling, B cell output can take place in neonatal mice lacking IL-7Ra until around two weeks of age (6). Furthermore, both the pre-BCR and mature BCR checkpoints are less stringent during neonatal life allowing for the positive selection of poly and self-reactive specificities in mice and men (7, 8, 9). This correlates with the output of self-reactive CD5+ B-1a cells in mice, which secrete natural antibodies important for the clearance of cellular debris and act as a first line of defense against pathogens (10). Despite these differences, the basis for the developmental switch in B cell output is not well understood.

One molecular program that distinguishes early life lymphopoiesis is centered around the post-transcriptional regulator LIN28B. Along with LIN28A, it is one of two mammalian homologs of the heterochronic *lin-28* RNA binding protein first identified in *Caenorhabditis elegans* (11). In the mammalian hematopoietic system, *Lin28b* expression is abundant during fetal life and gradually silenced within the first two to three weeks of life, critically controlling a neonatal switch in hematopoietic stem cell function and lineage output patterns (12, 13, 14). LIN28A/B plays an established role in regulating cellular growth and metabolism in various tissues, most famously through the direct inhibition of the biogenesis of the evolutionarily conserved Let-7 family of microRNAs. Its expression de-represses Let-7 targets which include the proto-oncogenes *Myc, Arid3a, Hmga2* and *Igf2bp1-3* (15, 16, 17). We and others have previously shown that ectopic LIN28A/B expression in the adult is sufficient to reinitiate a fetal-like gene expression signature and promote key aspects of fetal-like lymphopoiesis, including the efficient production of CD5+ B-1a cells (13, 14, 15, 18). These studies establish ectopic LIN28B expression as a powerful approach to understand the unique molecular program that governs early life B cell output. Furthermore, studies from both BCR poly-clonal and BCR transgenic mouse models make clear that LIN28B relaxes the central tolerance checkpoint to allow for the positive selection of self-reactive BCR specificities and thereby uniquely shapes the neonatal B cell repertoire (19–20). Immunophenotypically, this phenomenon is marked by the developmental progression of CD5+ Imm-B cells destined for the B-1a cell lineage (19). The mechanism for this effect is, although not fully understood, at least in part linked to augmentation of c-MYC protein levels in Imm-B cells (19) known to promote B cell positive selection (21).

A less understood mechanism of LIN28B action is through the direct binding of coding mRNAs to either positively or negatively regulate transcript stability and protein translation, in a context dependent manner (22, 23, 24, 25, 26, 27, 28, 29, 30, 31). In particular, several studies have demonstrated the ability of LIN28B to promote the translation of ribosomal proteins, through Let-7 independent direct binding to their transcripts in human ES cells (32), transformed human kidney (22) and neuroblastoma (31) cell lines. However, the role of the protein synthesis pathway in LIN28B dependent B cell output is not known. Here we demonstrate that LIN28B can directly interact with ribosomal protein transcripts in primary B cell precursors. Using an inducible mouse model we show that LIN28B expression during adult B cell development elevates protein synthesis during the small Pre-B and Imm-B cell stages. Finally, we show that elevated protein synthesis is a hallmark of neonatal B cell development that depends on endogenous *Lin28b* expression and critically supports the generation of CD5+ B-1a cells early in life. We conclude that LIN28B augments protein synthesis to potentiate the unique features of early life B cell output.

## Materials and methods

### Mice

tet-LIN28B mice were generated by intercrossing *Col1a^tetO-LIN28B^* (JAX: #023911) mice, carrying Doxycyclin (Dox) inducible Flag-tagged human *Lin28b* cDNA under the endogenous *Col1a* promoter, to *Rosa26^rtTA*m2^* (JAX: #006965) mice and were originally obtained from the laboratory of George Daley (Harvard Medical School) (33). Monoallelic transgene expressing mice were used for experiments. *Rosa26^rtTA*m2^* heterozygous littermates were used as control mice. For *in vivo* experiments, Doxycycline (Dox) chow (200 mg/kg, Ssniff, cat#A112D70203) was fed to tet-LIN28B and littermates for 10 days before analysis. B6.Cg-*Lin28b^tm1.1Gqda^*/J (Lin28b KO) (JAX: #023917), C57BLKS-*Rp124^Bst^*/J (Bst) (JAX: #000516) and Nur77-GFP (Jax #016617) were from the Jackson laboratory. Wildtype mice were from Taconic (B6NTAC, Taconic). The *Lin28b-eGFP* knock-in strain was generated as previously described (34). Adult mice were used at 10 to 16 weeks of age, neonates were analyzed at the indicated days post-birth. Male and female mice were used interchangeably throughout the experiments. Animal husbandry and experimental procedures were performed in accordance with ethical permits issued by the Swedish Board of Agriculture.

### Bone marrow cultures

Adult bone marrow B cell precursors from untreated adult mice were expanded and enriched *ex vivo*, as previously described (35). Briefly, single cell suspension of red blood cell lysed bone marrow cells from untreated adult mice were plated in complete RPMI-1640 (Fischer Scientific, cat#21875091) (supplemented with 10% FBS, HEPES, NEAA, Sodium Pyruvate, Penstrep, 2-mercaptoethanol) and incubated for 15 min at 37°C. The non-adherent cells were isolated and seeded at a cell density of 0.75-1.5 million/mL using complete RPMI-1640, IL-7 (20 ng/mL, Peprotech, cat#217-17), and Dox (0.1 μg/ml, Sigma-Aldrich, cat#D3072). Note that Dox treatment was always initiated *ex vivo* at the start of the culture. The cell culture medium was refreshed every second day.

### RNA Immunoprecipitation Sequencing (RIP-seq)

Cells from day 5 bone marrow cultures from tet-LIN28B or littermate control mice were used for lysis and RIP using Magna RIP RNA-Binding Protein Immunoprecipitation kit (Millipore, cat#17-700) according to the manufacturer’s instructions. Briefly, 10% of the total lysate was kept as input sample. 4 μg of anti-Flag antibody (Sigma) and 50 μl of A/G beads (Millipore, cat#16-663) were added to the lysate and incubated for 3 hours. Beads were washed 6 times with ice-cold RIP wash buffer (Millipore, cat#17-700). RNA was extracted using RNAzol (Sigma-Aldrich, cat#R4533) and Direct-zol RNA MicroPrep kit (Zymo Research, cat#R2060). For RNA quantification and quality control respectively, Qubit with RNA HS kit (Thermofisher Scientific, cat# Q32852) and Agilent Bioanalyzer with RNA Pico kit (Agilent, cat#5067-1513) were used according to the manufacturer’s instructions. 20ng of total RNA was used for the cDNA synthesis and amplification with the SMART-Seq v4 Ultra Low Input RNA Kit for Sequencing (Takara, cat#634889) according to instructions. Library tagmentation and indexing was performed using Nextera XT DNA Kit (Illumina, cat#FC-121-1030) and Nextera XT v2 set A Indexing kit (Illumina, cat# 15052163). Library quantification and library size were measured using Qubit DNA HS kit (Thermofisher Scientific, cat#Q32851) and BioAnalyzer DNA HS kit (Agilent, cat#5067-4626). The samples were run in NextSeq 550 using NextSeq High output 150 cycles 2×76bp.

The sequencing was aligned to mm10 using the STAR aligner (v2.7.1) and counted using RSEM (v1.28). After mapping, the RSEM result files were loaded into R and their FPKM values were used for the analysis. To filter out lowly expressed genes, the initial data (covering 54,752 genes) was filtered for genes that had at least 50 FPKM value in the input and IP of both tet-Lin28b samples (1,459 genes), which were then used for all the analysis presented in Figure 1. The individual sample FPKM values before and after filtering are included below.

| Sample | Initial reads (FPKM) | Filtered reads (FPKM) |
| --- | --- | --- |
| #1 tet-Lin28b input: | 1,320,000 | 1,090,000 |
| #1 tet-Lin28b IP: | 1,260,000 | 1,080,000 |
| #2 tet-Lin28b input: | 1,360,000 | 1,140,000 |
| #2 tet-Lin28b IP: | 1,230,000 | 1,040,000 |
| Ctrl input: | 1,380,000 | 1,140,000 |
| Ctrl IP: | 1,140,000 | 912,000 |
| Sum: | 7,690,000 | 6,402,000 |

**Figure 1.**
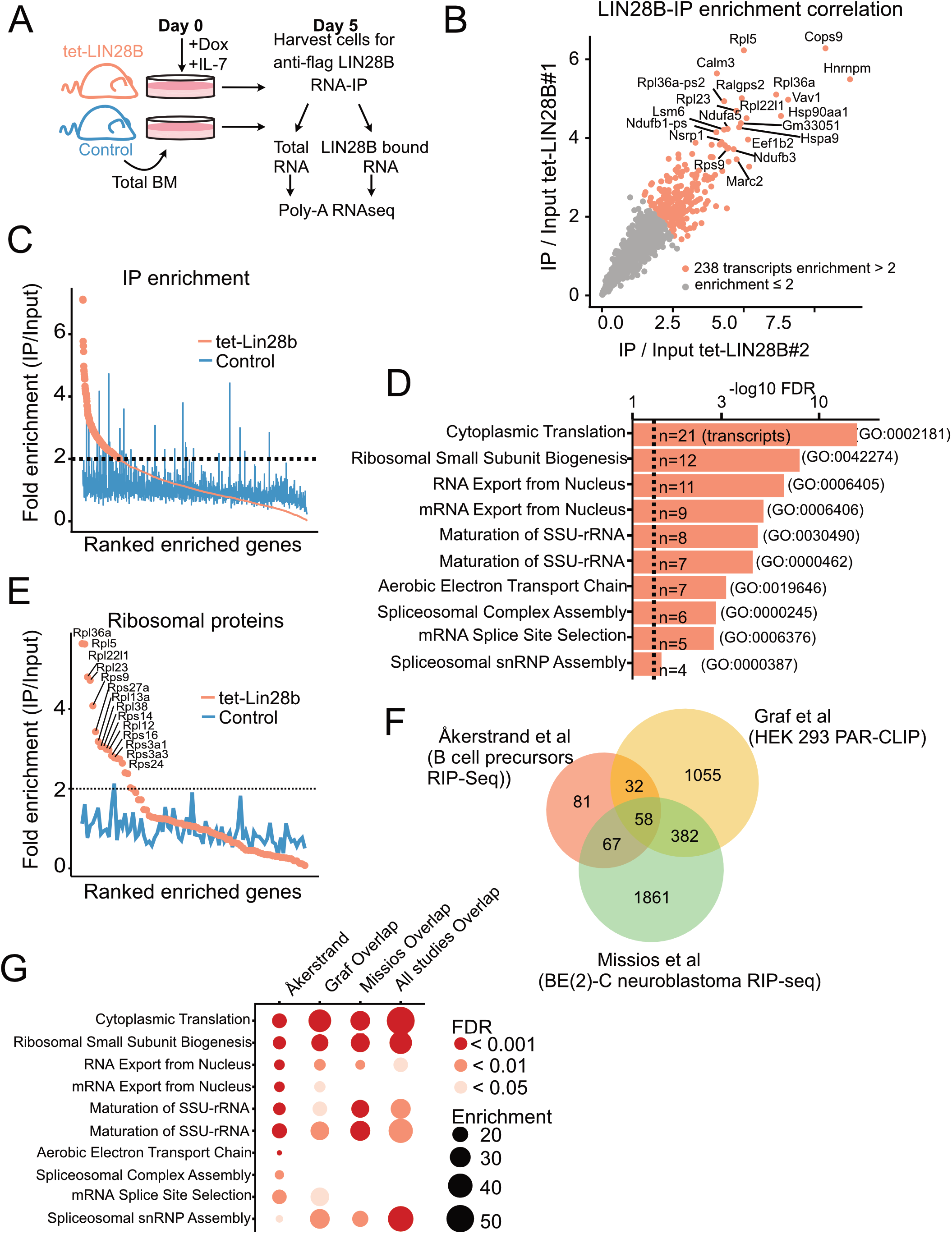
LIN28B binds to ribosomal protein transcripts in primary B cell precursors. **(A)** Schematic of experimental setup for the identification of LIN28B-bound mRNA targets in bone marrow (BM) B cell precursors. Uninduced adult total BM of the indicated genotypes were plated in IL-7 (20 ng/mL) and Doxycycline (Dox, 0.1 μg/mL) prior to RNA-IP for five days to allow for Pro-B cell expansion. Co-immunoprecipitated mRNA from two tet-LIN28B and one control mouse were individually subjected to Poly-A RNA-seq. **(B)** IP/input enrichment ratios from the two tet-LIN28B samples were plotted against each other showing high correlation between the two biological replicates. LIN28B IP/input enrichment ratio of greater than two-fold (n=238 transcripts) were used for subsequent analyses. **(C)** Average enrichment ratio ranking of the two tet-LIN28B samples compared to control. **(D)** Top ten gene ontology terms of the 238 enriched transcripts. **(E)** Graph showing the average enrichment ratio of all ribosomal protein coding transcripts ranked by IP/Input. **(F)** Analysis of overlapping genes between the 238 LIN28B enriched transcripts of this study (Åkerstrand et al) and previously published LIN28B-mRNA RIP-Seq and PAR-CLIP data sets (22–31). **(G)** Gene ontology analyses of overlapping interacting transcripts between the indicated studies.

Enrichment was calculated as IP over Input FPKM for each of the two tet-LIN28B samples, individually. 238 transcripts had an average enrichment of at least 2-fold and were considered for subsequent analyses. Littermate control sample was not considered when generating this list. Gene ontology analysis was done using the Panther database (pantherdb.org, release 17.0). Comparable results were obtained using the KEGG pathway and BioPlanet. All the sequencing data is accessible through GEO, GSE217788.

### Poly-A RNA sequencing

B cell precursor subsets were FACS sorted from adult BM of Dox induced mice of the indicated genotypes. RNA sequencing libraries were generated as previously described (19) and sequenced on Illumina NextSeq and NovaSeq. Note that FSC-A low, small Pro-B cells were excluded from the FACS sorting gate. All data was processed and analyzed as previously described (19). Data is accessible through GSE217788 except for Imm-B cells that was previously generated (19) and is accessible through GSE135603. GSEA used Wald’s stat as ranking metric. MSigDB Mouse collections Hallmark_MYC_target_v1.

### Metabolic labeling of nascent proteins

Cells were depleted of endogenous methionine by pre-culturing in methionine free RPMI (Fisher Scientific, cat#A1451701). After 45 minutes, the medium was supplemented with L-azidhomoalanine (AHA, 1 mM final concentration, Fisher Scientific, cat#C10102) and labeled for one hour. AHA uptake was visualized using Click-iT Plus Alexa Fluor 555 Picolyl Azide Toolkit (Fisher Scientific, cat#C10642), following the manufacturer’s protocol. Briefly, up to 3 x 10^6^ cells were fixed in 400 μL of 4% formaldehyde (Fischer Scientific, cat#28906) for 20 minutes at room temperature, washed, and then permeabilized in equal volume of 0.1% saponin (Sigma, cat#S4521) for 20 minutes. Alexa Fluor 488 Alkyne (Fisher Scientific, cat#A10267) was used in place of Alexa Fluor 555 for FACS analysis.

### EdU labeling for cell cycle analysis

Analysis was performed in accordance with the manufacturer’s instructions (Invitrogen, cat#C10634). Briefly, single cell suspension of red blood cell lysed bone marrow cells was plated in complete RPMI-1640 (Fischer Scientific, cat#21875091) and labeled with EdU (10 μM) for 30 minutes. Cells were then stained for surface proteins before fixation and permeabilization. The EdU was Click-iT ligated to Alexa Fluor 555 and analyzed with DAPI for separation according to cell cycle phase.

### Flow cytometry

Bone marrow cells were extracted by crushing bones from hind- and front limbs, hips, and sternum, using mortar and pestle. Peritoneal cavity was isolated by flushing adult mice with 8 mL of FACS buffer (Hank’s Balanced Salt Solution (Gibco, cat#14175-053) supplemented with 0.5% Bovine Serum Albumin (Sigma, cat#A9647) and 2 mM EDTA (Invitrogen, cat#15575-038)), or neonates 1-8 mL depending on the age. Red blood cell lysis was performed on bone marrow and spleens using ACK lysis buffer (Fischer Scientific, cat#A1049201), only for mice that were at least 19 days old. Lineage depletion was carried out before cell sorting by MACS Cell Separation and LS columns (Miltenyi Biotec, cat#130-042-401) to deplete CD3+, TER119+, or Gr-1+ cells, according to manufacturer’s instructions.

Antibody staining of surface antigen was performed using FACS buffer, at a cell density of 1 x 10^7^ cells per 100 μL of FACS buffer with antibodies, for 30 minutes at 4°C. Analysis of intracellular antigens was done by using the same fixation and permeabilization as was described under “Metabolic labeling of nascent proteins” or using BD Cytofix/Cytoperm fixation/permeabilization kit (BD Biosciences, cat#554714). Maximum of 3 x 10^6^ cells per reaction. All flow cytometry experiments were gated on live CD19+ B cells and the following immunophenotypes: Pro-B cells (CD93+IGM-cKIT+), Pre-B cells (CD93+IGM-cKIT- and FSC-A to separate large from small), Imm-B cells (CD93+IGM+), peritoneal cavity B-1a B cells (CD5+CD23–B220low and CD43+), and peritoneal cavity B-2 B cells (B220high CD23+ and CD43–). All flow cytometry experiments were performed at the Lund Stem Cell Center FACS Core Facility (Lund University, Lund, Sweden) using BD Fortessa, BD Fortessa X20, BD FACS Aria III, or BD FACS Aria II instruments. Cell sorting was performed using a 70 μm nozzle, 0.32.0 precision mask, and a flow rate below 6000 events/s. Analysis was performed using FlowJo version 10. Antibodies are listed in Supplemental table S1.

### Bone marrow HSPC transplantation

4000 bone marrow Lineage^-^ Sca1^+^ cKit^+^ (LSK) donor cells (CD45.1) from day 10 neonatal *Rpl24^WT/Bst^* or WT mice were FACS sorted and mixed at a 1:2 ratio with 8000 sorted CD45.1 competitor LSK cells and 200,000 CD45.1 total BM support cells. Cells were transplanted into lethally irradiated (2 × 450 cGy) and CD45.1+ CD45.2+ double positive adult recipients by tail vein injection. Endpoint analysis was performed 13–14 weeks after transplantation.

### Western Blot

Sorted B cells were lysed in RIPA buffer (CST, cat#9806S), protease inhibitor (Roche, cat#4693132001), and PMSF (CST, cat#8553S). Loading samples were prepared with Laemmli buffer (Bio-Rad, cat#1610747) with β-mercaptoethanol (Bio-Rad, cat#1610710). Proteins were separated by SDS–polyacrylamide gel electrophoresis (Any kD Mini-Protean TGX gel, Bio-Rad, cat#4568125) and transferred to a nitrocellulose membrane (Trans-Blot Turbo Mini Nitrocellulose Transfer Packs, Bio-Rad, cat#1704158). Membranes were blocked for 1 hour at RT in 5% skimmed milk. Antibodies are listed in Supplemental table S1. The ECL Select kit (GE Healthcare, cat#RPN2235) was used to detect protein bands with a ChemiDoc XRS+ system (Bio-Rad). Band intensity was quantified by Image Lab Software v6.1(BioRad). Each lane was automatically detected with manual adjustments to subtract the background and each band was individually selected followed by quantification of peak area of obtained histograms.

For RIPseq, 10% of the each indicated protein extract was collected to validate the immunoprecipitation efficiency. Proteins were separated by SDS–polyacrylamide gel electrophoresis (12% Mini-Protean TGX gel, Bio-Rad, cat#4561043) and transferred to a nitrocellulose membrane (Trans-Blot Turbo Mini Nitrocellulose Transfer Packs, Bio-Rad, cat#1704158). Membranes were blocked for 1 hour at RT in 5% skimmed milk. Antibodies are listed in Supplemental table S1.

### qRT-PCR

50.000 sorted cells were lysed in RNAzol (Sigma-Aldrich, cat#R4533) and total RNA was extracted using Direct-zol RNA MicroPrep kit (Zymo Research, cat#R2060) according to the manufacturer’s instructions. Total RNA (8 μl) was used in random primed cDNA synthesis with TaqMan Reverse Transcription Reagents (Applied Biosystem, cat#N8080234). Diluted cDNA 1:2 (4 μl) was used in every qPCR reaction using KAPA probe Fast qPCR kit (Kapa Biosystem, cat#KK4706) with primers targeting 45S (Thermofisher Scientific, cat# Mm03985792_s1) and 18S (Thermofisher Scientific, cat# 4333760T) rRNA transcripts. The relative expression levels of rRNA were evaluated using log2 (2ΔCt) calculation.

### Statistical analysis

All statistical analysis was carried out in R (version 4.1.2) and RStudio (version 1.4.1717), using functions from the following packages: tidyverse (version 1.3.1), DESeq2 (version 1.32.0), rstatix (version 0.7.0), fgsea (version 1.18.0), and ggpubr (version 0.4.0). Statistics for bulk RNA-seq was generated using the DESeq2 package without custom settings. Data from non-sequencing methods were imported into R and statistics calculated using Wilcox or T-test, as indicated by the figure legends. Asterisks indicate the P-value, *<0.05, **<0.01,***<0.001,****<0.0001.

## Results

### The LIN28B mRNA interactome in primary B cell precursors is enriched for ribosomal protein transcripts

Using an inducible tet-LIN28B mouse model expressing Flag-tagged human *Lin28b* cDNA upon doxycycline (Dox) induction (*Col1a^tetO-LIN28B^ Rosa26^rtTA*m2^*) (33), we and others have previously demonstrated that LIN28B expression is sufficient to re-initiate fetal-like hematopoietic output in adult life (14, 19, 36, 37). To better understand the underlying mechanism, we performed immunoprecipitation of its mRNA interactome in developing B cells. To obtain sufficient cell numbers for interactome profiling, adult B cell precursors from uninduced tet-LIN28B and control mice were expanded for 5 days by culturing whole bone marrow cells in the presence of IL-7 and supplemented with Dox *ex vivo* to induce LIN28B expression. After five days with IL-7, the cultures reached above 80% B cell purity (Supplemental figure 1A). Cells were harvested and subjected to anti-FLAG RNA immunoprecipitation (IP) coupled with poly-A selected RNA-sequencing (RIP-seq) (Figure 1A). Immunoprecipitation efficiency of LIN28B was validated by western blot analysis (Supplemental figure 1B). Total RNA sequencing of corresponding input samples was used for calculating RIP-seq enrichment. LIN28B RIP/input ratio of the technical replicates showed a high degree of correlation (R=0.93, p < 2.2e-16) (Figure 1B) and were averaged for downstream analyses. 238 transcripts displayed a higher than two-fold RIP/input enrichment in tet-LIN28B samples but showed no trend in the control (Supplemental Table S2 and Figure 1C). Gene ontology (GO) analysis revealed a prominent enrichment for ribosomal protein coding transcripts (Figure 1D, E), suggesting that ribosomal function might be a core target pathway of LIN28B during B cell development. Interestingly, similar observations were previously made in other cell types, including 293T cells (22) and BE(2)-C neuroblastoma cells (31). Indeed, cross comparison of our identified targets with these studies yielded cytoplasmatic translation as the predominant overlapping GO term (Figure 1F-G). Our analysis also identified other GO terms involved in RNA splicing and oxidative phosphorylation, consistent with previous LIN28A/B interactome studies in various cell types (23–38). We conclude that the direct LIN28B mRNA regulon in the context of primary bone marrow B cell progenitor cells is dominated by transcripts implicated in universal biosynthetic pathways, which are shared across a wide range of cell types. In particular, our findings implicate the ribosome as a potential target for LIN28B regulation in B cell progenitors.

### LIN28B enhances protein synthesis in the small Pre-B and Imm-B stages of B cell development

LIN28B bound mRNA targets have been linked to both translational increase and decrease in a context dependent manner (22, 24, 27, 31, 39, 40). In the case of ribosomal protein transcripts, enhanced protein translation was shown by SILAC (stable isotope labeling using amino acid in cell culture) metabolic labeling (22) as well as polysome profiling (31). To assess ribosomal protein levels during B cell development, we performed western blot analysis on sorted B cell precursors from tet-LIN28B and control mice following 10 days of Dox induction *in vivo* (Supplemental figure 2A). We selected RPL36, RPL23 and RPS9 from the RIP-seq enriched transcript list and found increased protein expression upon LIN28B induction in the Pro-B and large Pre-B subsets (Figure 2A). Differences were less apparent for the small Pre-B and Imm-B subsets (Supplemental figure 2B), which have lower cellular protein content. Overall, these results are consistent with enhanced translation of LIN28B bound ribosomal protein encoding transcripts.

**Figure 2.**
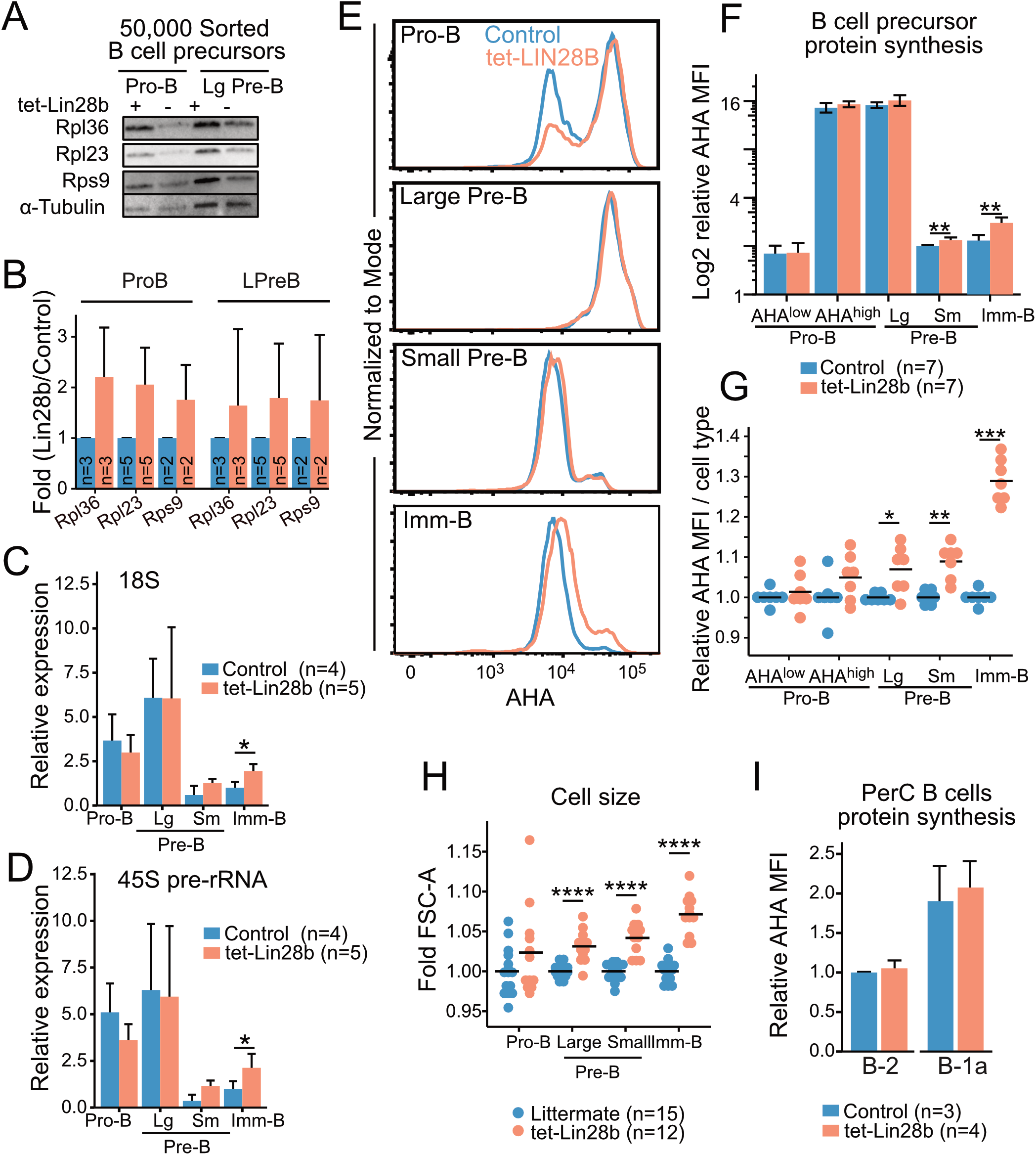
LIN28B enhances protein synthesis in small Pre-B and Imm-B cells. **(A)** Representative western blot analysis for the indicated ribosomal proteins from 50,000 FACS sorted Pro-B and large Pre-B cells from adult tet-LIN28B or littermate control mice following 10 days of *in vivo* Dox induction. **(B)** Quantification of western blot data from 2-5 biological replicates and three separate experiments (See materials and methods). **(C-D)** qRT-PCR for 18S rRNA or 45S pre-rRNA from 50,000 FACS sorted adult B cell progenitors. Expression values were normalized relative to the littermate control Imm-B cells. Large and small Pre-B cells are indicated as Lg and Sm respectively. **(E)** Representative FACS histogram overlays of L-Azidohomoalanine (AHA) uptake, corresponding to their protein synthesis rates, in the indicated subsets from adult mice. **(F-G)** AHA median fluorescence intensity (MFI) for indicated BM B cell precursor stage as shown in E. **(F)** Data is presented as relative values to one control small Pre-B cell subset for each experiment. **(G)** Data is normalized per cell type to one control sample for each experiment. **(H)** Measurement of FSC-A MFI. Data is normalized per cell type to one control sample for each experiment. **(I)** AHA MFI for the indicated, peritoneal cavity (PerC) B cell subsets, plotted relative to control B-2 values. Wilcox test was used to calculate p-values. *p<0.05, **p<0.01, ***p<0.001. Bars show mean, error bars show standard deviation.

Given this increase in ribosomal proteins, we expected a corresponding increase in overall ribosomal content upon tet-LIN28B induction. To this end, we measured the 18S rRNAs by qRT-PCR of FACS sorted B cell precursors. To our surprise, Pro and large-Pre-B cells did not exhibit any LIN28B dependent increase in 18S levels. Instead, ribosomal content first began to increase in a LIN28B dependent manner in the quiescent small Pre-B cell stage and reached statistical significance in Imm-B cells (Figure 2B). Additional analysis of the 45S precursor rRNA showed the same trend (Figure 2C). Thus, an increase in ribosome biogenesis takes place subsequent to the observed increase in ribosomal protein expression during LIN28B induced B cell development. To evaluate whether this observation correlated with changes in global protein synthesis rates, we measured newly synthesized proteins in freshly isolated B cell precursors using metabolic labelling with the methionine analog L-azidohomoalanine (AHA) (41). As expected, FACS readout of AHA uptake from Dox induced control and tet-LIN28B mice showed highly dynamic protein synthesis levels during B cell development that peaks in the large Pre-B cell subset (Figure 2D). Pro-B cells exhibited a bimodal profile with respect to protein synthesis that correlated with cell size (AHA^high^ or AHA^low^, Supplemental figure 2C). Although there was a tendency towards an increased frequency of AHA^high^ Pro-B cell in tet-LIN28B induced mice (Figure 2D, Supplemental figure 2C-D), the changes were not statistically significant. Increased global protein synthesis was observed in tet-LIN28B induced small Pre-B and Imm-B cell stages, thereby mirroring the observed effects on ribosome biogenesis. This was accompanied by increased cell size as measured by FACS (FSC-A) (Figure 2G), and a moderate increase in the fraction of proliferating cells within the Imm-B subset (Supplemental figure 2E). These findings are consistent with elevated protein synthesis boosting biomass accumulation and cell cycle entry during the later stages of B cell development. Interestingly, analysis of mature B cells showed that protein synthesis in peritoneal cavity B-1 and B-2 cells were not affected by ectopic LIN28B (Figure 2H). Taken together, we conclude that LIN28B enhances ribosome content and global protein synthesis levels specifically during the later stages of bone marrow B cell development.

### Lin28B augments protein synthesis during IL-7 independent B cell maturation

We sought to understand the stage dependent effects of LIN28B expression on protein synthesis by considering the signals that control dynamic changes in cellular metabolism and biosynthesis during B cell development. We have previously shown that Lin28B enhances the output of CD5+ Imm-B cells, and that CD5 levels correspond to BCR self-reactivity (19). To assess whether enhanced protein synthesis in Imm-B cells is linked to higher signaling levels through the newly selected BCR we compared CD5 expression with AHA uptake by tet-LIN28B induced Imm-B cells. Indeed, we observed a positive correlation between the two readouts suggesting that enhanced protein synthesis is dictated by BCR signaling strength (Figure 3A-B). To strengthen this interpretation, we confirmed that CD5+ Imm-B cells correspond to those with the highest level of Nurr77-GFP reporter expression (Figure 3C), a well-established measurement of antigen receptor stimulation in mice (42–43). Interestingly, Nur77-GFP was not altered in small Pre-B cells upon tet-LIN28B induction despite a ~10% increase in protein synthesis at this stage (Figure 2F). Taken together, our data show that LIN28B induced protein synthesis precedes the positive selection of CD5+ Imm-B cells and is in line with its permissive effect on the developmental progression from the small Pre-B to the Imm-B stages.

**Figure 3.**
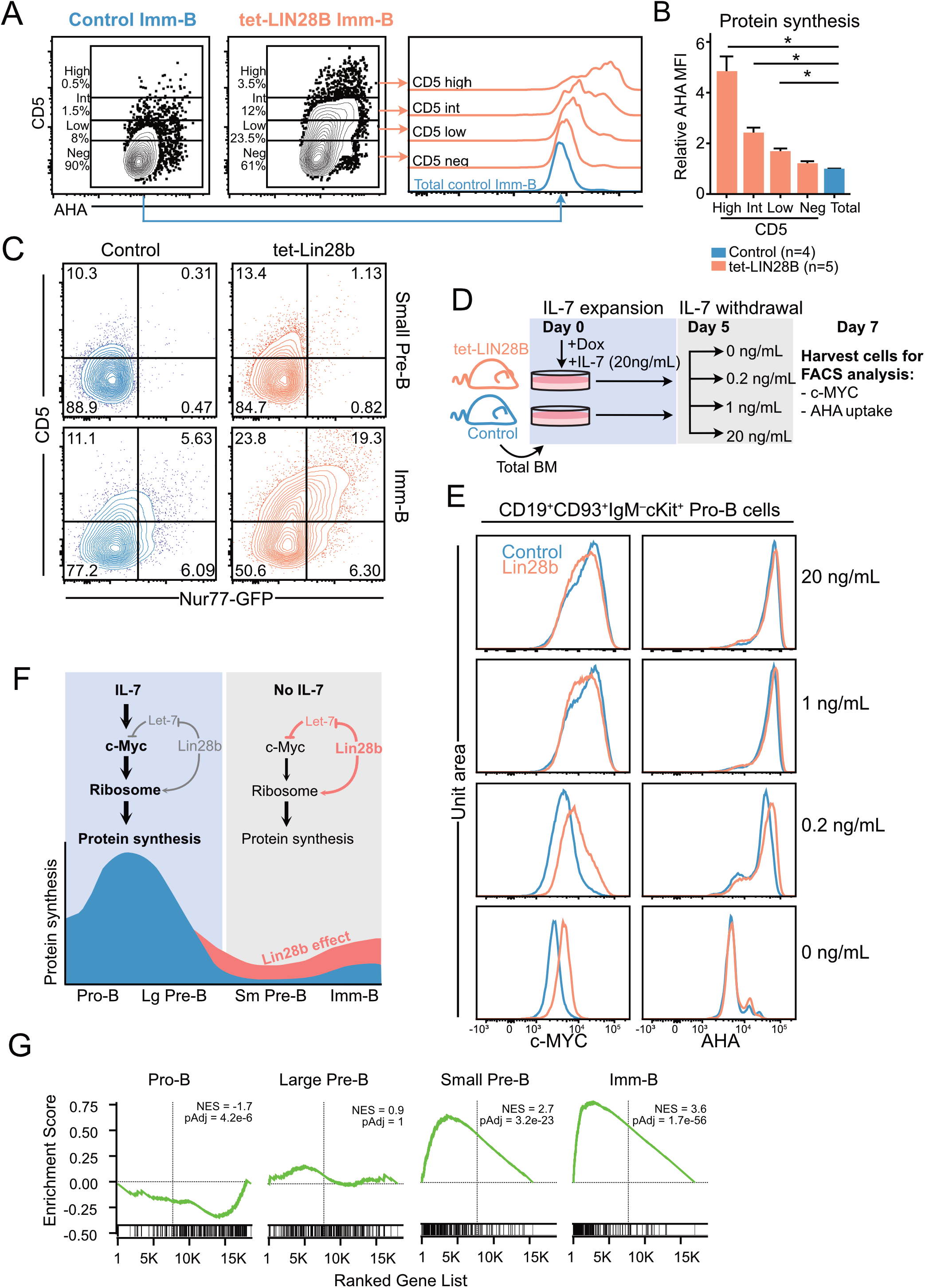
Interrogating the stage dependent effects of LIN28B on protein synthesis during B cell development. **(A)** (Left) Representative FACS plots showing CD5 expression in adult Imm-B cells from the indicated genotypes and (right) corresponding AHA uptake of the indicated subsets. **(B)** AHA MFI within individual CD5 slices, after normalization to AHA MFI in littermate control Imm-B cells. Wilcox test was used to calculate p-values in B. **p<0.01. Bars show mean ± standard deviation. **(C)** Representative FACS plot showing CD5 surface expression against Nur77-GFP levels for adult small Pre-B and Imm-B cells of the indicated genotypes. **(D)** Experimental setup for measuring the impact of IL-7 availability on sensitivity to LIN28B mediated effects on Pro-B cells. Following 5 days of expanding adult B cell precursors in 20 ng/mL IL-7, the cultures were split into conditions with decreasing concentrations of IL-7. Two days later, the cultures were analyzed by FACS for intracellular c-MYC and AHA uptake. **(E)** Representative FACS histograms for the indicated IL-7 conditions show intracellular staining for c-MYC (left) and AHA uptake (right) in Pro-B cells. **(F)** Schematic illustration of the proposed LIN28B mode of action on protein synthesis during B cell development. **(G)** Gene set enrichment analysis (GSEA) of differentially expressed genes between tet-LIN28B induced and control B cell precursor subsets as identified by RNAseq (See methods). Vertical line indicates Zero cross point. Enrichment for c-MYC target signature is shown.

The lack of LIN28B induced effects on protein synthesis in Pro-B cells was puzzling given the *Lin28b* dependent increase in ribosomal protein expression at this stage (Figure 2A). Considering that both IL-7 and LIN28B can act through elevating c-MYC expression, and that the c-MYC transcriptional program is a critical driver of cellular protein synthesis (2–44), we hypothesized that potent IL-7 signaling might mask any LIN28B mediated effects on the c-MYC / protein synthesis axis in Pro-B cells. To test this, we assessed LIN28B induced protein synthesis in Pro-B cells following IL-7 withdrawal *ex vivo*. B cell precursors were expanded from control and tet-LIN28B adult bone marrow under saturating IL-7 conditions for five days, before culturing under limiting doses of IL-7 (Figure 3D) as previously described (3). Two days following IL-7 withdrawal, cells were harvested and FACS analyzed for LIN28B-dependent effects on c-MYC expression and global protein synthesis. Interestingly, although overall c-MYC and protein synthesis levels of Pro-B cells decreased upon IL-7 withdrawal, the decrease was partially alleviated by tet-LIN28B induction (Figure 3E left). A similar trend was observed for protein synthesis (Figure 3E right) at limiting IL-7 concentrations. Thus, LIN28B has the capacity to enhance the c-MYC / protein synthesis axis in Pro-B cells, but only under limiting IL-7 availability. Taken together, our results provide a likely explanation to the stage dependent effects of LIN28B expression on protein synthesis during B cell development (Figure 3F). In Imm-B cells, enhanced protein synthesis is linked to the increased signaling strength of BCR specificities licensed by LIN28B expression. Prior to the Imm-B cell stage, effects on protein synthesis are uncoupled from the BCR, but is dictated by IL-7 responsiveness. During the Pro-B cell stage, potent IL-7 mediated protein synthesis masks the milder effects by LIN28B which only become visible upon entry into the small Pre-B cell stage whereby IL-7Ra expression is fully silenced. This scenario is supported by RNAseq data from sorted B cell precursor subsets from tet-LIN28B and control adult bone marrow, which showed a LIN28B induced increase in c-MYC target genes specifically upon entry into the small Pre-B cell stage (Figure 3G). Taken together, our results suggest a critical role for LIN28B in the small Pre-B cell stage as cells loose IL-7 responsiveness and point toward a key role for c-MYC in LIN28B induced protein synthesis.

### Elevated protein synthesis is a hallmark of neonatal B cell development

Endogenous LIN28B expression is developmentally restricted. FACS analysis of LIN28B-eGFP reporter mice (34) captured its window of expression as eGFP gradually diminished to background levels after postnatal day 10 in B cell precursors and postnatal day 19 in hematopoietic stem and progenitor cells (HSPCs) (Figure 4A). This time-window correlates with the output of B-1a cells which ceases between day 10 and day 19 during unperturbed hematopoiesis (45). During the same window, protein synthesis in neonatal bone marrow small Pre-B and Imm-B cells was elevated compared to the adult (Figure 4B). This was dependent on endogenous *Lin28b*, as 3-day-old KO neonates had a significantly lower rate of protein synthesis and decreased cell size during the small Pre-B and Imm-B cell stages (Figure 4C, D). These defects were primarily qualitative and not accompanied by changes in subset frequency (Figure 4E). We conclude that *Lin28b* dependent increases in protein synthesis during the small Pre-B and Imm-B cell stages represent a hallmark of early life B cell development.

**Figure 4.**
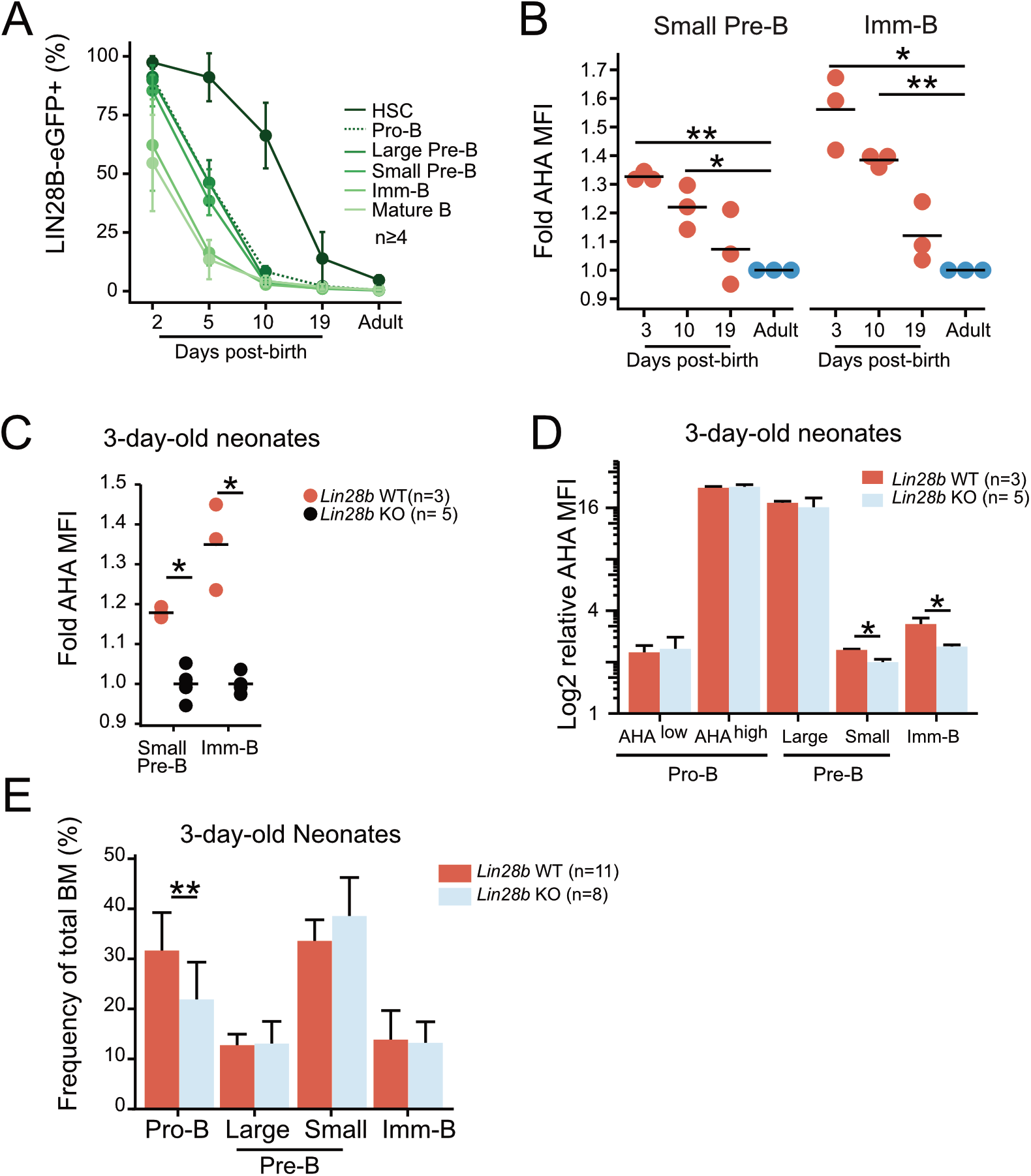
Elevated protein synthesis during neonatal B cell development is dependent on LIN28B. **(A**) Analysis of BM hematopoietic stem cells (HSCs: Lineage−, Sca1+, cKit+, CD150+, CD48-, FLT3+) and B cell precursors from *Lin28b-eGFP* mice at the indicated ages. The average frequency and standard deviation of *Lin28b*-eGFP positive cells is shown for each time point. **(B)** AHA MFI from *Lin28b*-WT neonates at the indicated ages are plotted relative to the uptake in WT adult control mice for each subset. **(C)** AHA MFI in 3-day-old neonates of the indicated genotypes. Values are normalized within each experiment to one *Lin28b* KO neonate per cell type. **(D)** Same as in C but normalized to one *Lin28b* KO small Pre-B cell sample within each experiment. **(E)** Frequency of BM B cell precursors in 3-day-old neonates. T-test was used to calculate p-values in B, Wilcox test was used to calculate p-values in C-E. **p<0.001, *p<0.05. Bars or line show mean, error bars show standard deviation.

### Neonatal B cell development relies on a heightened level of protein synthesis

To investigate the importance of elevated protein synthesis for neonatal B cell development, we took advantage of the ribosomal protein hypomorphic *Rpl24^WT/Bst^* mouse model. This model carries a mutation that disrupts *Rp124* mRNA splicing and protein expression (46). The resulting defect in ribosome biogenesis causes a moderate reduction in overall cellular protein synthesis which has no apparent impact on normal adult B cell development (41), but can diminish Eμ-MYC driven B lymphomagenesis known to be ‘addicted’ to heightened ribosomal function (47). Neonatal B cell development has not previously been examined in *Rp124^WT/Bst^* mice. Given our findings of higher protein synthesis rates, we hypothesized that neonatal B cell development would be more sensitive to constraints in protein synthesis capacity (Figure 5A). Indeed, analysis of *Rpl24^WT/Bst^* bone marrow revealed a decrease in the overall CD19+ B cell frequency on postnatal day 3 and day 10 that was resolved by day 19 (Figure 5B). This translated into a concomitant decrease in the frequency of B cell precursor subsets in the bone marrow of *Rp124^WT/Bst^* neonates (Figure 5C) and a time-limited decrease in the frequency of mature B cells in both the spleen (Figure 5D) and the peritoneal cavity of neonatal mice (Figure 5E). Thus, neonatal B cell development is more sensitive to constrained protein synthesis with signs of ribosome ‘addiction’. To verify that these effects are cell intrinsic during B cell development, we performed competitive transplantations of neonatal *Rpl24^WT/Bst^* or littermate bone marrow HSPCs into lethally irradiated congenic recipients (Figure 5F). Analysis of donor versus competitor reconstitution of recipient PerC B cells at 14 weeks post-transplantation showed a selective decrease in B-1a cells from *Rp124^WT/Bst^* donors. In contrast, contribution to the B-2 compartment, which relies on continuous HSPC dependent influx, was comparable between the *Rpl24^WT/Bst^* and wildtype donor HSPCs (Figure 5G). Taken together, these findings demonstrate that neonatal B cell output is selectively sensitive to reduced protein synthesis in a cell intrinsic manner.

**Figure 5.**
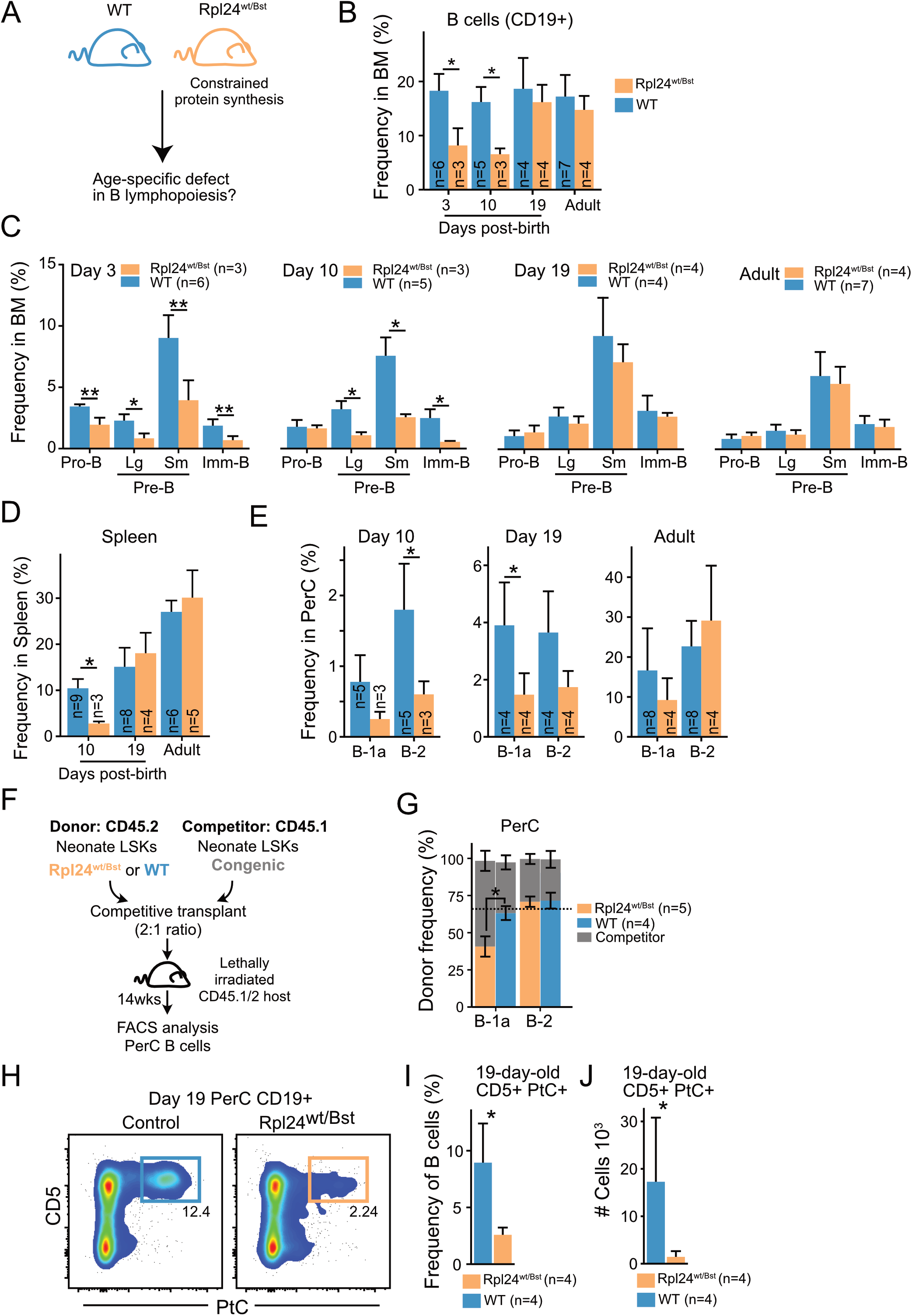
Neonatal B cell development relies on enhanced protein synthesis. **(A)** *Rp124^wt/Bst^* and wildtype littermates were analyzed by FACS at different ages for phenotypic differences during BM B cell development. **(B)** Frequency of CD19+ B cells out of total BM cells at the indicated ages. **(C)** BM B cell precursor subset frequencies at the indicated ages presented as percent of total BM cells. **(D)** Frequency of splenic CD19+ B cells out of total splenocytes at the indicated ages. **(E)** Mature PerC B-1a and B-2 cell frequencies out of total PerC cells at the indicated ages. **(F)** Experimental overview for assessment of cell intrinsic defect of the *Rp124^wt/Bst^* mutation. Lineage−, Sca1+, cKit+ (LSK) hematopoietic stem and progenitor cells (HSPCs) from 10-day-old donor and competitor mice of the indicated genotypes were sorted and transplanted into lethally irradiated recipients. 13-14 weeks later, the steady state donor and competitor derived mature PerC B cells were analyzed to assess B-1a and B-2 cell output. **(G)** CD19+ PerC B cells of the indicated subsets were compared for donor versus competitor contribution. The expected frequency based on donor:competitor ratio is indicated by the dashed line. **(H)** Representative FACS plots showing the frequency of CD5+ PtC-liposome reactive B cells in 19-day old littermates. **(I)** CD5+PtC+ frequency out of PerC CD19+ B cells in 19-day-old mice. **(J)** Absolute number of CD5+PtC+ B cells in PerC in 19-day-old mice. Wilcox test was used to calculate p-values. *p<0.05, **p<0.01, ***p<0.001. Bars show mean ± standard deviation.

The B-1a cell subset is enriched for the VH11 and VH12 encoded reactivity against the self-membrane component phosphatidyl choline (PtC) known to originate early in life. To investigate the qualitative impact of constrained protein synthesis on B-1a cell composition, we analyzed the frequency and absolute number of PtC reactive B-1a cells in 19-day-old Rp124^*WT/Bst*^ mice using fluorescently labelled PtC liposomes. This timepoint was selected to minimize influence by peripheral homeostatic expansion (48). Indeed, FACS analysis revealed a dramatic decrease in the frequency and absolute number of PtC reactive B cells in the peritoneal cavity of *Rpl24^WT/Bst^* neonates (Figure 4H-J). We therefore conclude that elevated protein synthesis is a central distinguishing parameter of neonatal B lymphopoiesis that potentiates the output of CD5+ B-1a cells carrying self-reactive BCRs.

## Discussion

In this study, we identified a role for cellular protein synthesis in regulating the unique aspects of early-life B cell development including the time restricted output of self-reactive B-1a cells. Elevated protein synthesis was a defining feature of neonatal B cell development that was mediated at least in part by the endogenous expression of the heterochronic RNA-binding protein LIN28B. As the expression of LIN28B naturally waned during the first weeks of life, protein synthesis became reduced, and ectopic LIN28B expression in the adult was sufficient to recapitulate elevated protein synthesis small Pre- and Imm-B cells. Consistent with previous reports in other cell types, we showed that LIN28B directly bound to ribosomal protein transcripts in primary B cell precursors, correlating with their enhanced protein expression. Furthermore, an increase in ribosomal biogenesis and protein synthesis coincided with LIN28B induced c-MYC expression in the quiescent small Pre- and Imm-B cell stages. Finally, using *Rp124^WT/Bst^* mice as a model for constrained protein synthesis capacity, we showed that neonatal B cell development relied on heightened protein synthesis levels – a hallmark of elevated c-MYC activity. The precise extent and mechanism by which LIN28B cooperates with c-MYC to promote cellular protein synthesis was not disentangled in this study. One likely possibility is that LIN28B may post-transcriptionally act on a c-MYC induced transcriptome to facilitate downstream translation, biomass accumulation and B cell developmental progression.

Protein synthesis is the most energy consuming biosynthetic process in the cell and subject to the constraints imposed by hundreds of components required for ribosome assembly (49). Indeed, protein synthesis rates give an accurate account of the overall cellular metabolism (50). And yet, its ability to control the development and function of immune cells that require rapid expansion and contraction of cellular growth has been an under-explored subject of investigation. Our results using AHA metabolic labelling to measure protein synthesis confirmed dynamic changes in cellular metabolism that underly proliferative bursts following successful BCR checkpoints (1). Importantly, we demonstrated the ability of LIN28B to partially alleviate the biggest drop in protein synthesis upon entry into the small Pre-B cell stage. The restricted metabolism of small Pre-B cells was previously implicated as a mechanism to safeguard the central tolerance checkpoint through enforcing nutrient stress and negative selection upon strong BCR engagement (5). This notion is highly compatible with our observation that LIN28B promoted protein synthesis to augment the output of self-reactive CD5+ B-1 cells early in life. The latter is a hallmark of early life B lymphopoiesis and offers an attractive explanation to the long-known prevalence of self-reactive specificities in neonatal mice and men (7–51). Thus, we propose that the relatively mild increase in protein synthesis mediated by LIN28B provides sufficient relaxation of the metabolic constraints during B cell maturation to qualitatively alter the B cell output.

In Pro-B cells, IL-7 signaling and downstream mTORC1 activation is the principal driver of c-MYC expression and acts to promote biomass accumulation and proliferation (2). In this study, we showed that both IL-7 signaling and LIN28B expression act on the c-MYC / protein synthesis axis to promote developmental progression during B cell development and that the powerful effects of IL-7 renders LIN28B function redundant in Pro-B cells. This finding is relevant in the context of understanding the ontogenic differences in IL-7 dependency during B lymphopoiesis. It well-established that the strict requirement for IL-7 mediated signaling, which is characteristic of adult murine B lymphopoiesis, does not apply during fetal and neonatal life (6). We have previously demonstrated that LIN28B over-expression during adult hematopoiesis is sufficient to phenocopy fetal-like behavior by circumventing strict IL-7 dependency (13). Here, our findings that LIN28B partially cushions the impact of IL-7 withdrawal through promoting the shared downstream c-MYC / protein synthesis axis may offer an explanation to the lessened reliance on IL-7 during early life B lymphopoiesis (6).

The RIP-seq experiment presented in this study led us to identify changes in protein synthesis levels as a distinguishing parameter between neonatal and adult B cell development. However, the experiment has several limitations. First, the LIN28B interactome in IL-7 expanded bone marrow Pro-B cell cultures may not exactly reflect that in primary small Pre-B cells. Unfortunately, the amount of material needed for RIPseq is not feasible to obtain from quiescent small Pre-B cells, and data extrapolation should be made with caution. Second, the low number of biological replicates of this experiment prevents statistical analysis and limits the number of enriched transcripts identified. Nevertheless, since our data did detect the same top gene ontology terms as previous datasets from other tissues, they serve as confirmatory evidence in the context of B cell precursors. A recent LIN28B interactome study in hematopoietic progenitors expressing the oncogenic fusion protein MLL-ENL identified both Rps9 and Rpl23 transcripts as direct targets of LIN28B, offering further support to our findings in the hematopoietic system (40). Despite these limitations, our approach resulted in the identification of a relevant regulatory axis for early-life B lymphopoiesis. In the context of neonatal immunity, we and others have previously demonstrated that neonatal antigen exposure produces unique B cell memory clones not generated by adult mice immunized with the same antigen (45–52). Our current findings that enhanced protein synthesis quantitatively and qualitatively alters neonatal B cell output implicates ribosomal control in the formation of a unique pre-immune repertoire in neonatal mice and has important implications for the layered formation of a complex adaptive immune system (53, 54, 55).

## Acknowledgements

We thank Jonas Ungerbäck and Johanna Tingvall-Gustafsson for help with bioinformatics. We thank Jenny Hansson, Leal Oburoglu, and Svetlana Soboleva for their input. Finally, we also thank the FACS and Bioinformatics core facilities at the Lund Stem Cell Center for their support.

## Funding

J.Y. was supported by the Swedish Research Council, the Swedish Cancer Society, the European Research Council (715313), Knut and Alice Wallenberg Foundation and the Wenner-Gren Foundations.

---

**Table S1.** Antibodies and reagents used for Flow Cytometry, Western Blot, or IP.

| Antigen | Fluorochrome | Clone | Manufacturer | Reference# |
| --- | --- | --- | --- | --- |
| 7-AAD | - | - | BD Bioscience | 559925 |
| B220 | APC Cy7 | RA3-6B2 | BioLegend | 103224 |
| B220 | PE CF594 | RA3-6B2 | BD Biosciences | 562313 |
| CD117 | BV421 | 2B8 | BD Biosciences | 562609 |
| CD117 | PE | 2B8 | Invitrogen | 12-1171-82 |
| CD19 | APC Cy7 | 1D3 | BD Biosciences | 557655 |
| CD19 | BV786 | 1D3 | BD Biosciences | 563333 |
| CD23 | PE Cy7 | B3B4 | BioLegend | 101614 |
| CD25 | PE eFluor610 | PC61.5 | Invitrogen | 61-0251-82 |
| CD3 | PE Cy5 | 145-2C11 | BioLegend | 100310 |
| CD43 | FITC | S7 | BD Biosciences | 553270 |
| CD5 | BV421 | 53-3.3 | BD Biosciences | 562739 |
| CD5 | PE | 53-7.3 | BioLegend | 100608 |
| CD93 | APC | AA4.1 | BioLegend | 136510 |
| CD93 | BV786 | AA4.1 | BD Biosciences | 740941 |
| c-MYC | Unconjugated | Y69 | Abcam | Ab32072 |
| Goat anti-Rabbit IgG (H+L) Cross-Adsorbed Secondary Antibody | AF647 | Polyclonal | Invitrogen | A-21244 |
| Gr-1 | PE Cy5 | RB6-8C5 | BioLegend | 108410 |
| IgM | FITC | 11/41 | BD Biosciences | 553437 |
| IgM | PE Cy7 | RMM-1 | BioLegend | 406514 |
| Alkyne, 5-isomer | AF488 | - | Invitrogen | A10267 |
| PtC | Marina Blue | DOPC/CHOL Liposomes | Formumax | F60103F-MB |
| Ter119 | PE Cy5 | TER-119 | BioLegend | 116210 |
| Tubulin alpha |  |  | CST | 2125 |
| RPL23 |  |  | Abcam | ab264369 |
| RPL36a |  |  | Invitrogen | PA5-70719 |
| RPS9 |  |  | Abcam | ab157125 |
| FLAG M2 |  |  | Sigma | F1804 |
| LIN28B |  |  | CST | 4196 |

## Notes

### Competing Interest Statement

The authors have declared no competing interest.

### Summary of Updates

Major reworking of text and figures

