## Supplemental Figure 1 for "Enhanced protein synthesis is a defining requirement for neonatal B cell development"

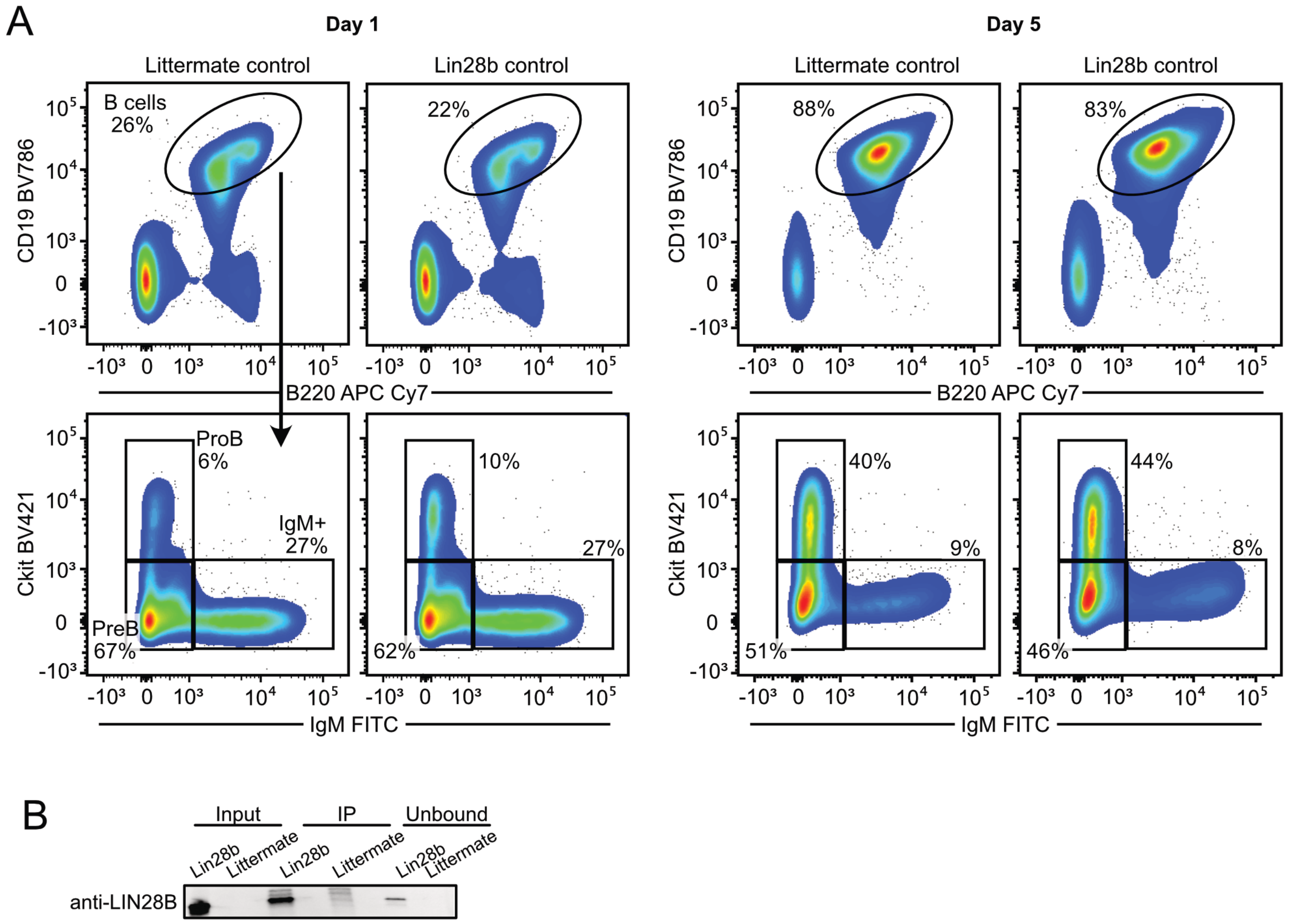


**Supplementary Figure 1.**

**(A)** Representative FACS plots of IL-7 bone marrow cultures on the indicated days. Whole bone marrow cells were cultured in presence of IL-7 (20 ng/mL) and doxycycline (0.1 µg/mL). **(B)** Western blot analysis for validating the IP of LIN28B.
