## Supplemental Figure 2 for "Enhanced protein synthesis is a defining requirement for neonatal B cell development"

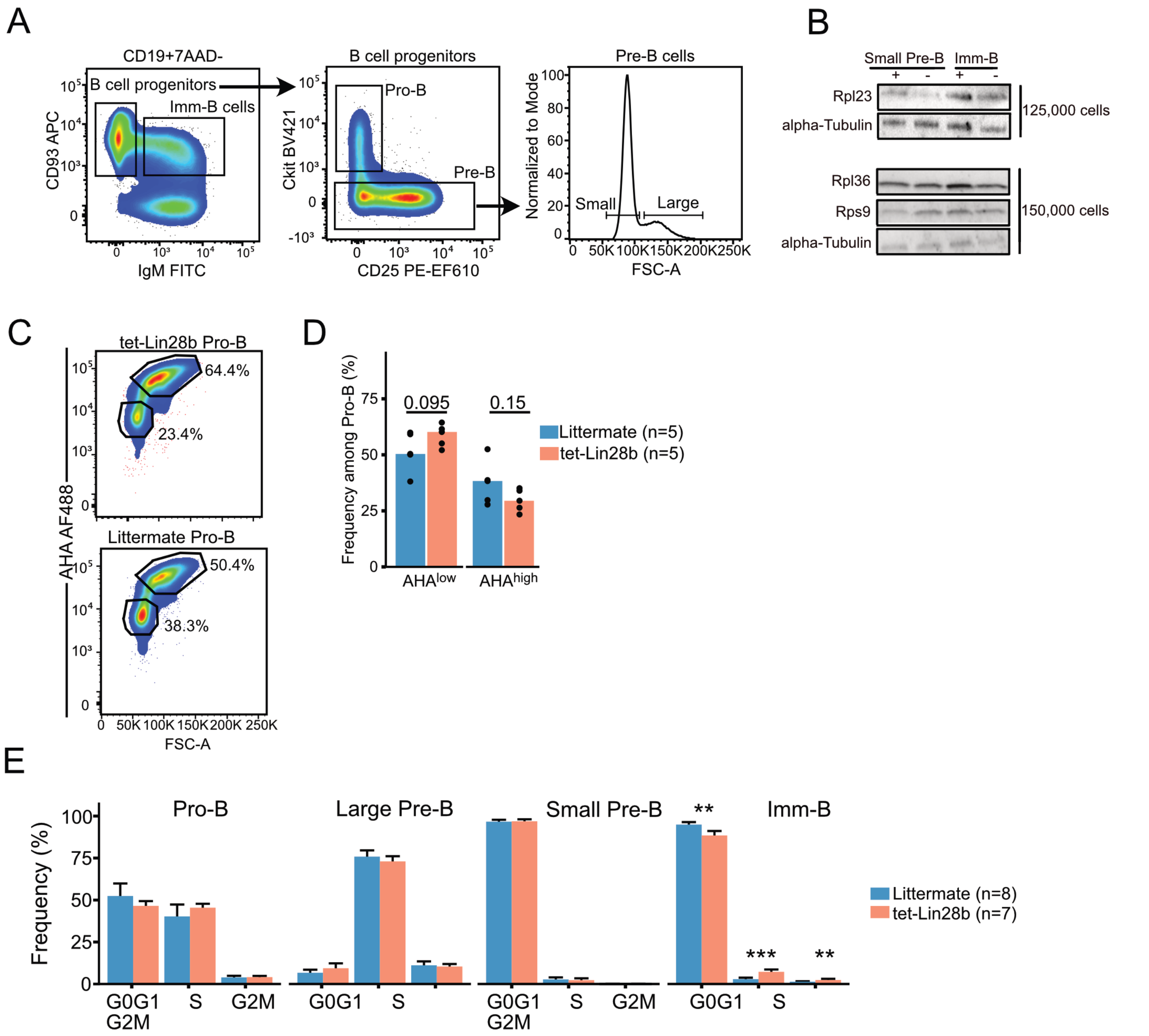


### Supplementary Figure 2.

**(A)** Gating strategy used for FACS sorting of B cell progenitors. **(B)** Representative western blot analysis for the indicated ribosomal proteins from sorted small Pre-B and Imm-B cells from adult tet-LIN28B or littermate control mice following 10 days of *in vivo* Dox induction. **(C)** Representative FACS plots showing the frequency of $AHA^{high}$ and $AHA^{low}$ cells within the Pro-B compartments from the indicated genotypes. **(D)** Summary of frequencies as shown in B. Data from three separate experiments. **(E)** Summary of cell cycle distributions as measured using EdU and 7AAD by FACS. Data from three separate experiments. Wilcox test was used to calculate p-values in B,D, and E. Bars show the mean ± standard deviation.
